# Human bone marrow organoids for disease modelling, discovery and validation of therapeutic targets in hematological malignancies

**DOI:** 10.1101/2022.03.14.483815

**Authors:** Abdullah O. Khan, Michela Colombo, Jasmeet S. Reyat, Guanlin Wang, Antonio Rodriguez-Romera, Wei Xiong Wen, Lauren Murphy, Beata Grygielska, Chris Mahoney, Andrew Stone, Adam Croft, David Bassett, Gowsihan Poologasundarampillai, Anindita Roy, Sarah Gooding, Julie Rayes, Kellie R Machlus, Bethan Psaila

## Abstract

A lack of models that recapitulate the complexity of human bone marrow has hampered mechanistic studies of normal and malignant hematopoiesis and the validation of novel therapies. Here, we describe a step-wise, directed-differentiation protocol in which organoids are generated from iPSCs committed to mesenchymal, endothelial and hematopoietic lineages. These 3-dimensional structures capture key features of human bone marrow - stroma, lumen-forming sinusoidal vessels and myeloid cells including pro-platelet forming megakaryocytes. The organoids supported the engraftment and survival of cells from patients with blood malignancies, including cancer types notoriously difficult to maintain *ex vivo*. Fibrosis of the organoid occurred following TGFβ stimulation and engraftment with myelofibrosis but not healthy donor-derived cells, validating this platform as a powerful tool for studies of malignant cells and their interactions within a human bone marrow-like milieu. This enabling technology is likely to accelerate discovery and prioritization of novel targets for bone marrow disorders and blood cancers.

**Significance Statement:** We present a 3D, vascularised human bone marrow organoid that supports growth of primary cells from patients with myeloid and lymphoid blood cancers. This model allows for mechanistic studies of blood cancers in the context of their microenvironment, and provides a much-needed*, ex vivo* tool for prioritization of new therapeutics.

## Introduction

The specialized bone marrow microenvironment maintains and regulates hematopoiesis, enabling adequate supply of blood cells to meet changing physiological requirements throughout life. Perturbations in the bone marrow hematopoietic niche contribute to the initiation and propagation of hematological malignancies. In addition, the stromal remodelling that occurs as a consequence of blood cancers contributes to bone marrow failure(1–4). Modelling bone marrow dysfunction is challenging, particularly in the context of human diseases. *In vitro* studies are limited to two dimensional (2D) systems and simple co-cultures, in which the relevant cell types are absent, and many human diseases are inadequately reproduced by mouse models. Patient-derived xenografts have been used to model disease and validate targets *in vivo*, but some malignancies and hematological cell subtypes do not engraft well, even when humanised murine models are used (5–11).

Advances have been made in modelling certain marrow components on ‘biochips’ (12–16), but the lack of specialized stroma, vascularization and active blood cell generation remains a limitation with these methods. Improved *in vitro* systems are required to enable more detailed mechanistic studies of human hematopoiesis and to allow functional interrogation of the pathways and crosstalk that drive bone marrow malignancies. The development and application of organoids – self-organizing, 3D, living multi-lineage structures – has the potential to facilitate translational research by enabling genetic screens and pharmacological modulation of disease pathobiology (17). Our goal was to generate a vascularised human bone marrow-like organoid that contains the key hematopoietic niche elements and supports active endogenous hematopoiesis, as well as the growth and survival of hematopoietic cells from adult donors including malignant cell types that are difficult to grow and study *ex vivo*. Such a system would offer a scalable and highly manipulable human model for mechanistic studies and drug development and, importantly, would reduce dependence on animal models.

To achieve this, we optimised a protocol in which human induced pluripotent stem cells (iPSCs) generate mesenchymal elements, myeloid cells and ‘sinusoidal-like’ vasculature in a format that resembles the cellular, molecular and spatial architecture of myelopoietic bone marrow. We confirmed the homology of these organoids to human bone marrow using multi-modal imaging approaches and single cell RNA-sequencing (scRNAseq). Crucially, in addition to modelling physiological hematopoietic cell-niche interactions, we showed that the organoids supported engraftment and survival of healthy and malignant hematopoietic cells from adult human donors, and enabled the screening of inhibitors of bone marrow fibrosis, a complication that occurs in patients with certain blood cancers and is associated with poor prognosis.

This addresses a long-standing need for 3D human bone marrow models for translational research where both niche and hematopoietic components are species and cell context specific, and creates a dynamic platform for high-throughput drug screening and studies aiming to understand disease pathways.

## RESULTS

### Mixed-matrix hydrogels containing Matrigel and type I and IV collagen are optimal for production of vascularised, myelopoietic organoids

To mimic the central bone marrow space (Fig. 1A), we devised a four-stage workflow to generate mesenchymal, vascular and myelopoietic marrow components. Human iPSCs were allowed to form non-adherent mesodermal aggregates (Phase I, days 0 – 3, Fig. 1B) before commitment to vascular and hematopoietic lineages (Phase II, days 3 – 5, Fig. 1B). The resulting cell aggregates were then embedded in mixed collagen-Matrigel hydrogels to induce vascular sprouting (Phase III, days 5 – 12, Fig. 1B). At day 12, sprouts were collected individually and cultured to form organoids in 96-well ultra-low attachment (ULA) plates (Phase IV, days 12 onwards, Fig. 1B).

**Figure 1:**
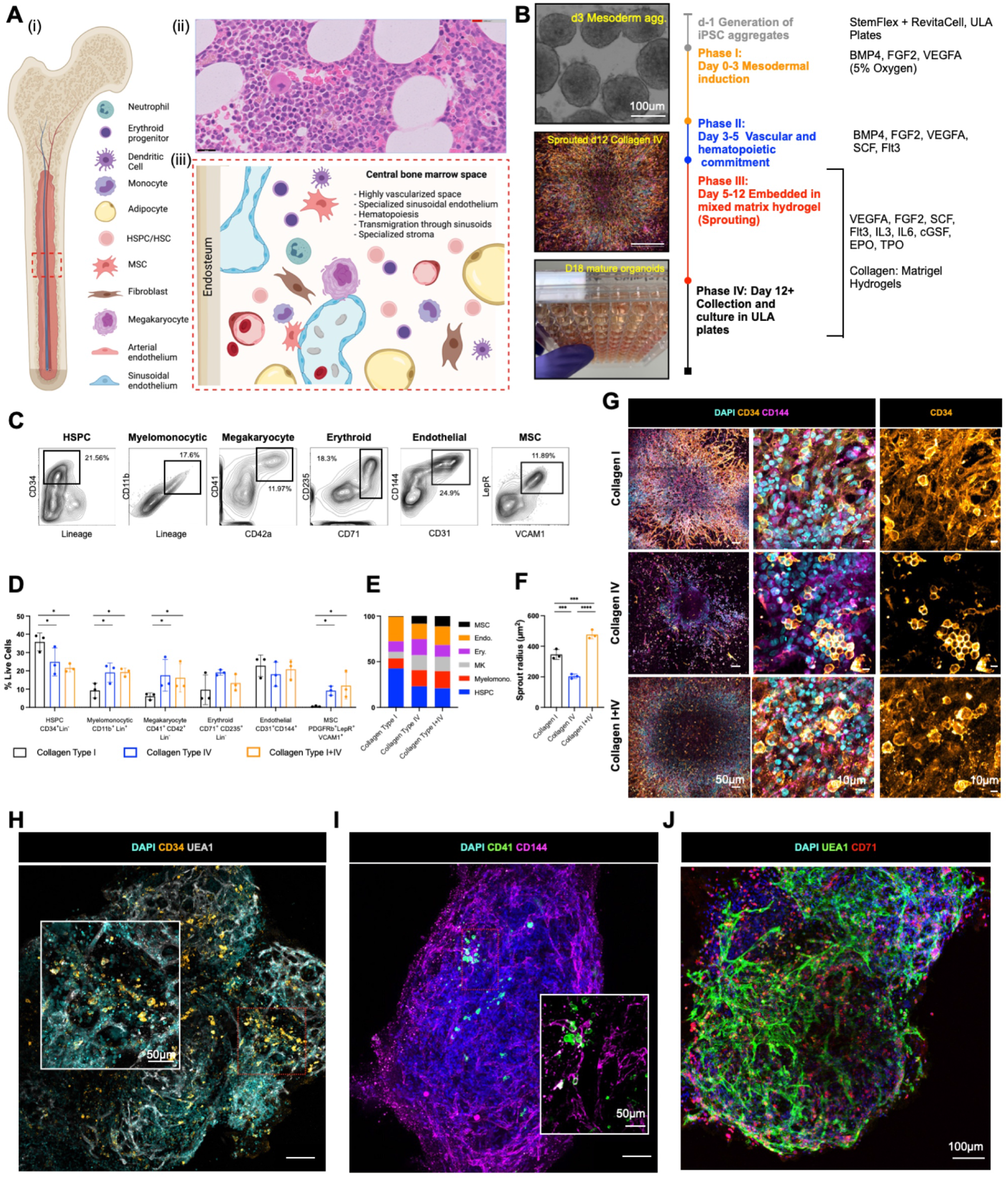
Mixed-matrix hydrogels containing Matrigel and type I and IV collagen are optimal for production of vascularised, myelopoietic organoids. **(A)** (i) Central bone marrow is a complex tissue including stromal, endothelial, hematopoietic stem and progenitor cells (HSPCs), myeloid and lymphoid subsets, (ii) H&E image of human bone marrow comprised of (iii) diverse hematopoietic and stromal cell types. (B) Differentiation workflow, in which iPSC aggregates underwent mesodermal induction (day 0-3) and commitment to hematopoietic and vascular lineages (day 3 - 5). Cell aggregates were then embedded in mixed matrix hydrogels comprised of Matrigel and collagen I, collagen IV, or collagen l+IV mix at a 40:60 ratio to support vascular sprouting. Key media components are listed for each phase. (C) Gating strategy and (D) quantification of stromal and hematopoietic cell types in day 18 organoids supported by Matrigel + collagen type I only, collagen IV only and collagen l+IV hydrogels. (E) Distribution of lineages as fractions of the whole organoid population. **(F)** Radius of endothelial sprouts and **(G)** sprouting day 12 organoids immunostained for nuclei (DAPI), CD34 and CD144 (VE-cadherin). **(G, H, I)** Whole organoid Z-stack imaging showing CD34^+^ HSPCs **(H)** CD41^+^ megakaryocytes **(I)** and **(J)** CD71^+^ erythroid cells dispersed throughout the organoids and associating with CD144+ UEA1+ vasculature. Schematic A created by Biorender.com. *p< 0.05, ** p < 0.01, *** p < 0.001, One-Way ANOVA with multiple comparisons (Fisher’s LSD) *n = 3* for endothelial sprout radius measurements, Two-Way ANOVA with multiple comparisons (Fisher’s LSD); *n* = 3 (3 independent differentiations, 15 pooled organoids each) for FACS Analysis. Representative images shown. **See also Figure S1**.

Hydrogels comprised of Matrigel plus type I collagen have previously been used to support the formation of iPSC-derived blood vessel organoids (18). However, type IV collagen is more permissive than type I collagen for myeloid and megakaryocyte maturation in standard 2D *in vitro* culture systems(19,20). We therefore compared hydrogels containing type I and/or type IV collagen plus Matrigel for the generation of stromal, endothelial and myeloid lineages. Distinct immunophenotypic hematopoietic stem and progenitor cell (HSPC, CD34^+^ Lin^-^) myelomonocytic (CD34^-^ CD11b^+^ Lin^+^), megakaryocyte (CD34^-^ Lin^-^ CD41 ^+^ CD42b^+^), endothelial (CD31^+^ CD144^+^), erythroid (CD34^-^ Lin^-^ CD71^+^ CD235a^+^) and MSC populations (CD31^-^ CD140b^+^ VCAM1^+^ LepR^+^) were detected when cells were digested at day 18 of differentiation (Fig. 1C – 1E). Organoids developed in collagen type I-only hydrogels contained the highest fraction of HSPCs but a low proportion of myelomonocytic cells, megakaryocytes and no MSCs, while collagen type IV and collagen types I+IV hydrogel-derived organoids yielded significantly larger myeloid and MSC populations, indicating that the addition of collagen IV created more favourable conditions for multi-lineage differentiation (Fig.1D & 1E, Supp. Fig. 1).

The hematopoietic niche contains a dense network of sinusoidal vessels, and the specialized sinusoidal endothelium regulates stem cell self-renewal and differentiation, progenitor maturation and platelet generation (21–25). To assessed vascularization of the organoids, we measured the degree of endothelial sprouting (26–29). Vascular sprout radii were significantly smaller in collagen type IV-only hydrogels compared to collagen types I-only and I+IV (average radius of 203.6 ± 14.61 μm *vs*. 345.9 ± 32.11 μm *vs*. 476.3 ± 28.82 μm respectively; Fig. 1F), and the density of CD34+ CD144 (VE-cadherin)-positive sprouts was significantly lower in collagen IV-only hydrogels (Fig. 1G). To assess vascularization and cellular architecture of the whole organoid in 3D, organoids were mounted in agarose and cleared with Ethyl Cinnamate for confocal Z-stacked imaging. This revealed an elaborate network of UAE1+ CD144+ vessels throughout the collagen I+IV-derived organoids (Figs. 1H – 1J). In addition to CD34^+^ HSPCs (Fig. 1H), CD41^+^ megakaryocytes (Fig. 1I) and CD71^+^ erythroid cells (Fig. 1J) were distributed throughout the organoid bodies and closely associated with the endothelium, as occurs in native bone marrow.

### Addition of VEGFC induced specialization of organoid vasculature to a bone marrow sinusoid-like phenotype

Having determined that collagen I+IV Matrigel hydrogels enabled multilineage differentiation, we sought to determine the optimal balance of endothelial growth factor support to generate organoid vasculature that resembles bone marrow sinusoids (23,24). Vascular endothelial growth factor (VEGF) A is a key regulator of blood vessel formation in health and disease, acting via the VEGF receptors VEGFR1 and 2 (30). Bone marrow sinusoidal endothelial cells express VEGFR1 and 2 as well as VEGFR3 (31), and VEGFC – the main ligand for VEGFR3 – was recently demonstrated to maintain the perivascular hematopoietic niche in murine bone marrow (32). We therefore tested the effect of adding VEGFC to the vascular sprouting phase (Day 5, Fig. 2A). Addition of VEGFC significantly increased expression of *FLT4* (encoding VEGFR3) in organoids, as well as HSPC adhesion molecules (*VCAM1, ITGA4*), HSPC-supporting growth factors and chemotactic cytokines (*CXCR4, FGF4;* Fig. 2B). VEGFA + C supplementation also induced retention of CD34 expression on organoid vessels, similar to native adult bone marrow vessels, while organoids stimulated with VEGFA alone expressed CD34 during vessel sprouting (days 5 - 12) but lost CD34 expression by day 18 (Fig. 2C).

**Figure 2:**
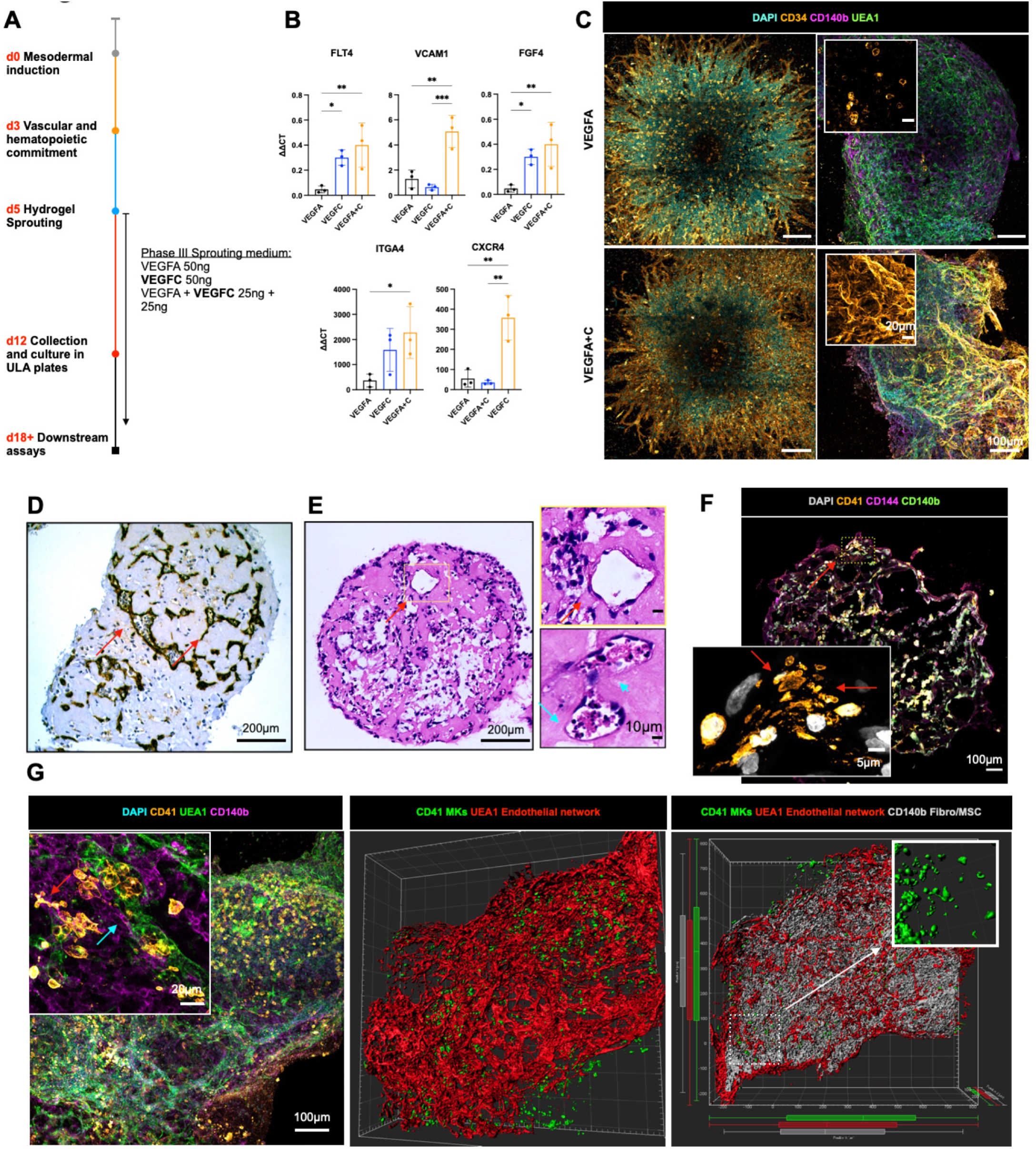
Addition of VEGFC induces specialization of organoid vasculature to a bone marrow sinusoid-like phenotype. **(A)** Organoids were supplemented with either VEGFA, VEGFC or VEGFA+C in the sprouting phase of differentiation (D5) in hydrogels. **(B)** mRNA expression of canonical cell surface receptors, growth factors and adhesion markers of bone marrow sinusoidal endothelium in VEGFA, VEGFC and VEGFA + C treated samples. ΔΔCt values relative to housekeeping (GAPDH) shown. Each data point represents 15 organoids pooled from independent differentiations. (C) Sprouting vessels positive for CD34 at day 12 in both VEGFA and VEGFA+C conditions **(D)** At day 18, vessels were CD34 negative in VEGFA-only treated organoids. **(E)** Immunohistochemical staining for CD34 and **(F)** Hematoxylin and Eosin staining of formalin-fixed, paraffin-embedded VEGFA+C organoid sections showed lumen-forming vessels (red arrows) containing hematopoietic cells (blue arrows) **(G)** Confocal imaging and 3D render of whole-mount VEGFA + C organoids showing CD41^+^ megakaryocytes (red arrow) closely associating with UEA1^+^ vessel network that is invested with CD140b^+^ fibroblast/MSCs (blue arrow) (left & centre image). 3D rendered megakaryocytes display proplatelet formation (right image, inset). (H) CD41^+^ megakaryocytes extending pro-platelet protrusions into vessel lumen. (* p < 0.05, ** p < 0.01, *** p < 0.001*, N = 3*, One-Way ANOVA with multiple comparisons [Fisher’s LSD]).

Organoid vessels formed clear lumens, containing extravasating hematopoietic cells (Fig. 2D & 2E). These vessels were invested with perivascular CD140b^+^ fibroblasts/MSCs (Fig. 2F), with CD41 ^+^ megakaryocytes closely associating with the vessels and extending proplatelet protrusions within the vessel lumen (Fig. 2F & 2G), with remarkable similarity to previously published *in vivo* images of thrombopoiesis occurring in calvarial bone marrow (33). Together, these data indicate that addition of VEGFC improved vascularization of the organoids and hematopoietic support.

### scRNAseq confirmed that hematopoietic and stromal cell lineages within organoids have transcriptional similarity to human hematopoietic tissues

To compare the cell types and molecular profiles of the organoids to human hematopoietic tissues in further detail, scRNAseq was performed on 13,449 cells from 3 independent organoid differentiations. After quality control (see Methods), 9,409 cells were included in downstream analyses. Distinct populations of the key hematopoietic and stromal cell subtypes were identified including HSPCs, erythroid, neutrophil, monocyte, megakaryocyte, eosinophil/basophil/mast (EBM), fibroblasts, endothelial cells and MSCs (Fig. 3A), annotated using Gene Set Enrichment Analysis with a curated list of 64 published gene sets (Fig. 3B, Suppl. Table 1) (34,35) as well as expression of canonical marker genes (Fig. 3C, Suppl. Table 2). Erythroid, megakaryocytic, monocytic/neutrophil and EBM populations demonstrated expression of *GYPA* and *KLF1, PF4* and *PPBP, CD14* and *RUNX1*, and *TPSB2* and *KIT* respectively (Fig. 3C). Within the stromal cell subsets, expression of *COL3A1*, platelet-derived growth factors (*PDGFRA/B*) and the key HSPC niche factor *CXCL12* (SDF-1) were expressed by fibroblasts and MSCs, with particularly high expression of *CXCL12* in the MSCs, suggesting a key role for MSCs in homing and maintenance of HSPCs to perivascular regions (Figs. 2F, 2G, 3C and Suppl. Fig. 2A & 2B). High expression of *PECAM1, CDH5* and *ENG* was detected in endothelial cells, confirming vascular specification (Fig. 3C, Supp. Fig 2A & 2B).

**Figure 3:**
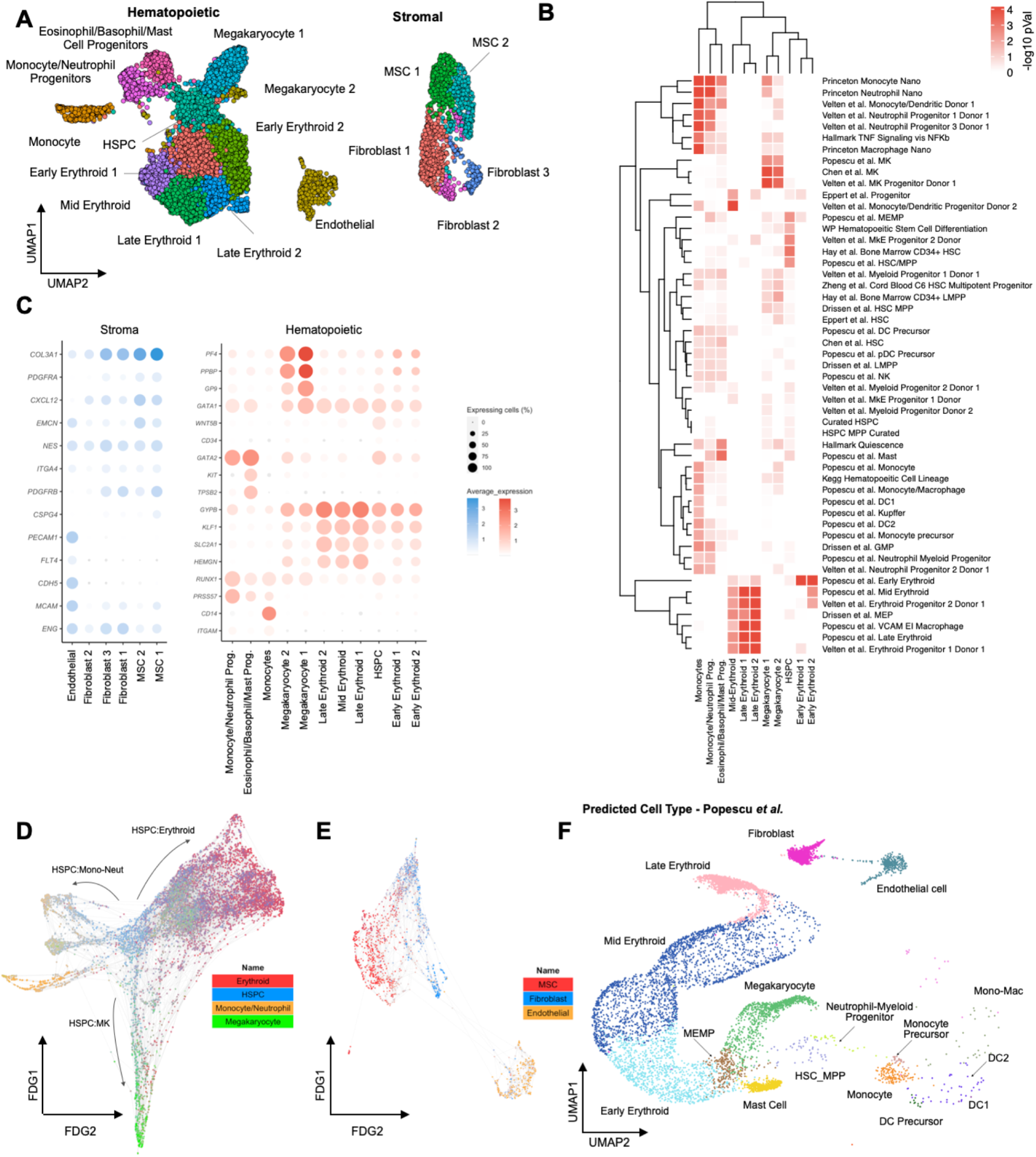
ScRNAseq confirmed that hematopoietic and stromal cell lineages within organoids showed transcriptional similarity to human hematopoietic tissues. **(A)** Uniform Manifold Approximation and Projection (UMAP) plot showing annotated cell clusters. **(B)** Gene Set Enrichment Analysis (GSEA) of differentially expressed genes for each cluster using a curated set of hematopoietic lineage gene sets. (C) Expression of canonical stromal and hematopoietic cell genes for each of the annotated clusters. Color scale represents the average level of expression and circle size shows % of cells within each cluster in which expression was detected for each gene. **(D & E)** Force-Directed Graph showing differentiation trajectories for **(D)** hematopoietic and **(E)** stromal compartments, superimposed with expression scores of lineage signature gene sets. (F) Organoid cells projected onto a published dataset of human hematopoietic and stromal cells using Symphony. **See also Figures S2 & S3 and Suppl. Tables 1 and 2**.

Trajectory analysis using a force-directed graph (FDG) showed that the organoids had generated cells that recapitulated the three main routes of hematopoietic myeloid differentiation (Fig. 3D & Suppl. Fig. 2C), similar to observations in human bone marrow (35). FDG analysis of the stromal cell populations showed independent routes of differentiation for endothelium and MSC/fibroblasts, as expected (Fig. 3E, Suppl. Fig. 2D).

To determine the transcriptional similarity of the bone marrow organoid cell types to native human hematopoietic tissues, the organoid scRNA-seq data was projected onto relevant human hematopoiesis single cell datasets of cells isolated from adult bone marrow (35,36), fetal liver and fetal bone marrow (37,38)using the Symphony package (39). This revealed extensive overlap of organoid-derived cells with HSPCs, myeloid subsets, fibroblast/MSC and endothelial cell types with the predicted cell types matching the cluster annotations (Fig. 3F,Suppl. Fig. 3), confirming that the iPSC organoid-derived cells had substantial homology to native human hematopoietic tissues.

### Bone marrow organoids recapitulate cellular and molecular cross-talk between hematopoietic, endothelial and stromal cells

To investigate the cellular and molecular interactions between hematopoietic, endothelial and stromal cell subtypes within the organoids, we mapped the expression of interacting receptor and ligand pairs across clusters. Complex communication networks were detected, both within and between hematopoietic and stromal cell compartments (Fig. 4A). Strong autocrine and paracrine interactions were predicted between MSCs, fibroblasts, endothelial cells, and monocytes, HSPCs and megakaryocytes, while erythroid cells showed weak interactions (Fig. 4A, Supp. Fig. 4A – C). Numerous interacting receptor-ligand partners were detected between megakaryocytes and endothelial cells (Suppl. Fig 4B, Suppl. Fig 5A), and megakaryocytes with MSCs (Suppl. Fig. 5B), indicating bi-directional regulatory interactions between these cell types. These included *NOTCH1-JAG1/2*, *FLT4-PDGFC*, *ANGPT2-TEK* and *FLT1-VEGFB* for endothelium:megakaryocytes, and *NOTCH1-DLL4*, *KIT-KITLG*, *FGF2-CD44*, *SELP-CD34* and *VEGFRA-FLT1* for megakaryocytes:endothelial cells (Suppl. Fig. 4B and Supp. Fig. 5A). Interactions between megakaryocytes and MSCs were dominated by growth factors produced by megakaryocytes, including transforming growth factor beta (TGFβ), platelet-derived growth factor (*PDGF*), fibroblast growth factor (*FGF*) and *VEGF* family members and their cognate receptors (Suppl. Fig. 5B). Similarly, monocytes and endothelial cells demonstrated abundant interacting partners indicative of regulatory interactions, including *TNF-NOTCH1*, *JAG1/2-NOTCH*, *SIRPA-CD47*, and *LGALS9*, *ICAM*, *VEGF* family members with cognate receptors (Suppl. Fig. 4C and Suppl. Fig. 5C). Significantly interacting partners between monocytes and MSCs/fibroblasts included *CXCL12-CXCR4*, *ICAM1-aXb2*, *SPP1-CD44*, interleukins 1 and 16, hepatocyte and fibroblast growth factors with their respective binding partners (Suppl. Fig 5D).

**Figure 4:**
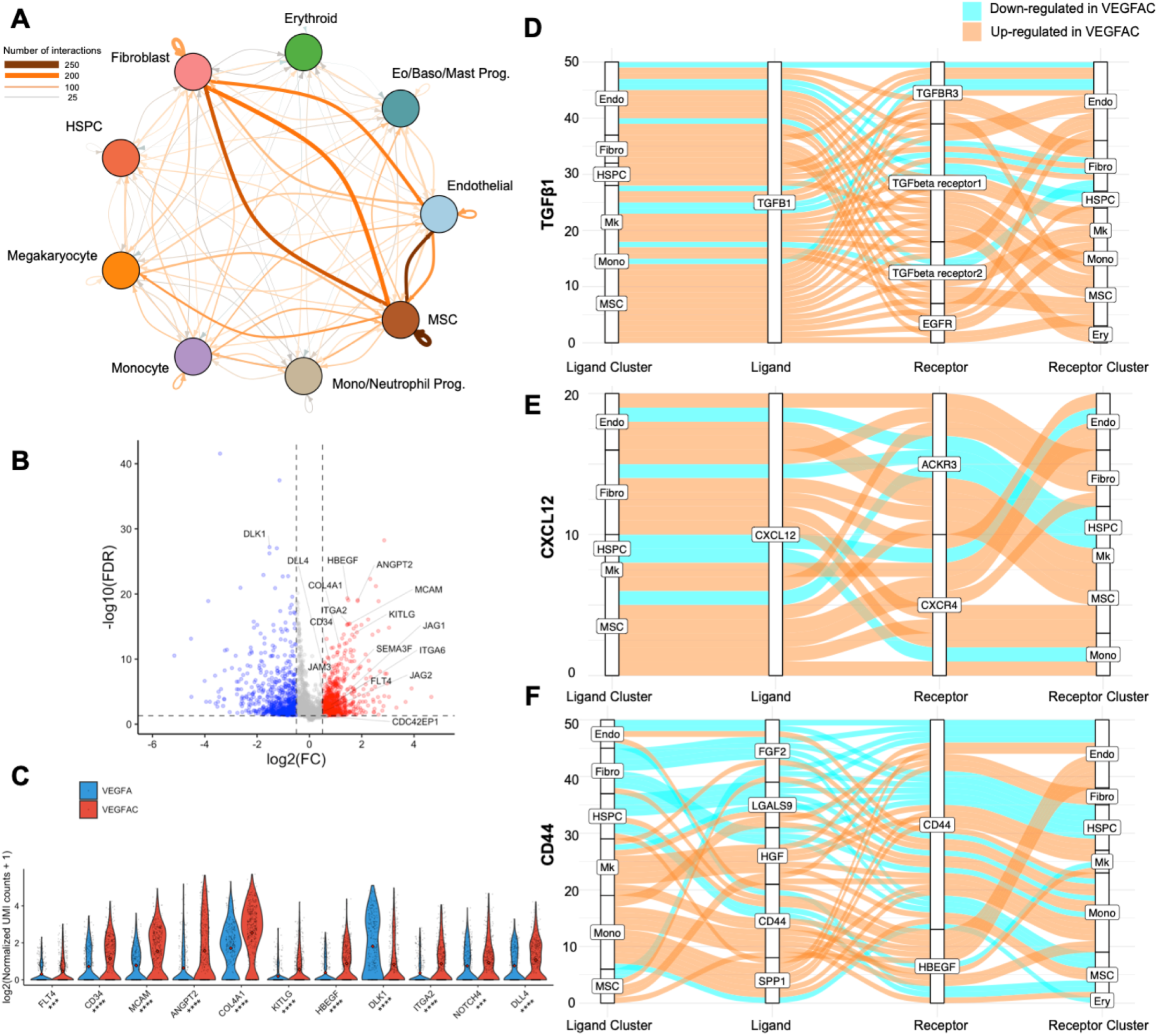
Extensive cellular and molecular interactions between hematopoietic, endothelial and stromal cells within organoids, with increased hematopoietic support from VEGFA+C stimulated vasculature. **(A)** Number of predicted interactions between cellular subsets within VEGFA+C-stimulated organoids. (B) Volcano plot and (C) violin plot showing differentially expressed genes between endothelial cells of VEGFA vs. VEGFA+C-stimulated organoids, with key hematopoietic support and sinusoidal endothelium genes highlighted. *P* values for comparison indicated below x-axis labels. **(D, E, F)** Sankey plots comparing **(D)** TGFβ, **(E)** CXCL12 and **(F)** CD44-mediated interactions in VEGFA+C vs. VEGFA stimulated organoids. (* p < 0.01, ** p < 0.05, *** p < 0.001, **** p < 0.0001 for pairwise comparisons, Wilcox test applied [FDR]). **See also Figures S4 & S5**.

Although a high number of regulatory interactions between hematopoietic and stromal compartments was detected, interactions between the stromal cell subsets (endothelial cells:MSCs:fibroblasts) were particularly strong (Fig. 4A). Significantly interacting partners identified included key regulatory molecules such as *JAG-NOTCH, VEGF, DLL4-NOTCH3, PDGF-PDGFR, ANPT1/2-TEK, IL33-IL33R, FGF* and *TGFB* (Supp. Fig. 5E) (40–42).

To explore the impact of addition of VEGFC to VEGFA in the differentiation protocol on the transcriptional phenotype of organoid vasculature, we compared the transcriptomes of endothelial cells captured from resulting organoids by scRNAseq. 2733 differentially-expressed genes were detected between endothelial cells generated with VEGFA only *vs*. VEGFA+C (Fig. 4B). Canonical markers of bone marrow sinusoidal endothelium were more highly expressed in endothelial cells from VEGFA + C cultures compared to VEGFA alone, including *FLT4* (VEGFR4), *CD34*, *MCAM*, *ANGPT2*, *COL4A1*, *COL4A2*, *ITGA2*, *CDC42EP1* and the notch ligand *DLL4* (24,37,43,44), while *DLK1* – a negative regulator of hematopoiesis (45) – was significantly downregulated in VEGFA + C-stimulated organoids (Figs. 4B & 4C).

In addition to improved hematopoietic support from endothelial cells, key regulatory axes were also upregulated across other cellular subsets in VEGFA+C organoids (Fig. 4D - F). TGFβ1 signalling primarily from MSCs, megakaryocytes and fibroblasts was overall increased in VEGFA+C organoids, across the different TGFβ receptors (Fig. 4D). Similarly, CXCL12 signalling via CXCR4 and ACKR3 receptors across both stromal and haematopoietic cell types was also overall increased (Fig. 4E), while the impact of VEGFC on signalling between CD44 and its binding partners was more mixed (Fig. 4F).

Together, these data indicate that communication networks between hematopoietic, mesenchymal and vascular compartments of VEGFRA+C-stimulated organoids reproduce many of the key regulatory interactions occurring within adult bone marrow (46–49).

### Bone marrow organoids model the TGFβ-induced bone marrow fibrosis that occurs in hematological cancers and provide an *ex vivo* platform for inhibitor screening

Pathological hematopoietic niche remodelling occurs in the majority of hematological malignancies. In certain cancers, particularly myeloproliferative neoplasms, myelodysplasia, acute leukemia and mast cell neoplasms, bone marrow fibrosis is a major cause of bone marrow failure, morbidity and associated with a poor prognosis (50,51). Fibrosis results from the excess production and release of pro-fibrotic cytokines, in particular TGFβ by hematopoietic cells, leading to the deposition of reticulin and collagen fibres by marrow stroma(50–53). To investigate whether the organoids could model pathological bone marrow fibrosis, we treated organoids with varying doses of TGFβ, which resulted in a dose-dependent increase in the expression of hallmarks of fibrosis, including collagen 1 and alpha smooth muscle actin (aSMA) (Fig. 5A), a markers of fibroblast activation(53). A significant increase in soluble IL11 was also observed (54)(Fig. 5B). Collagen deposition within the organoids was markedly increased following TGFβ treatment (Fig. 5C), with pronounced reticulin fibrosis (Fig. 5D), recapitulating changes seen in the bone marrow of patients with myelofibrosis.

**Figure 5:**
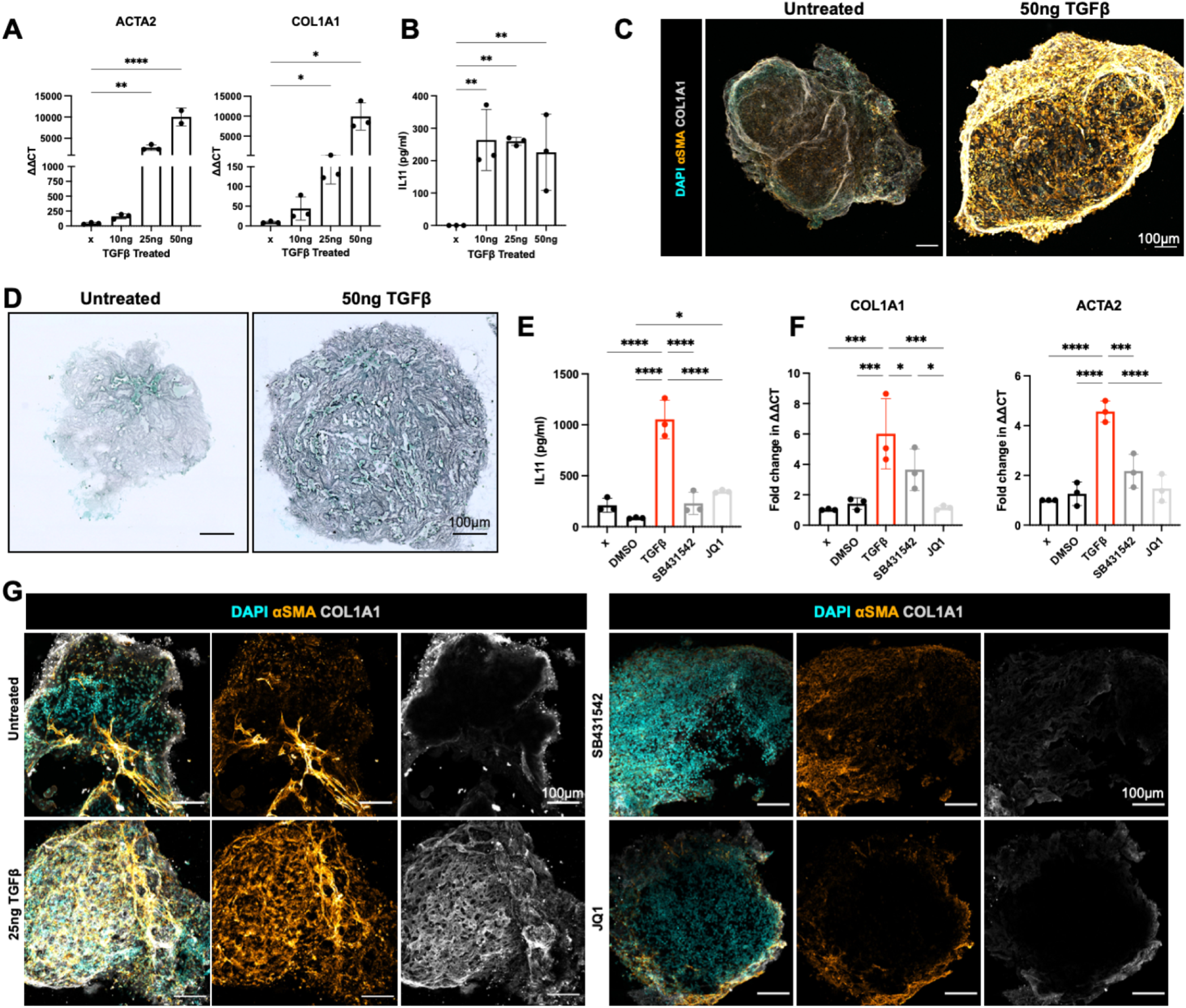
Bone marrow organoids model TGFβ-induced bone marrow fibrosis and enable inhibitor screening. **(A)** Organoids were treated with 10, 25, or 5Ong/mL recombinant TGFβ and evaluated by qRT-PCR for expression *cri ACTA2* (αSMA) and *COL1A1*, indicators of fibrosis. (B) Soluble IL11 detected in organoid media following treatment of organoids with TGFβ. (C) Confocal Z-stack images of whole, untreated and TGFβ (50ng/mL)-treated organoids stained for αSMA and COL1A1. (D) Reticulin staining in formalin-fixed, paraffin-embedded sections of TGFβ-treated organoids vs. control. **(E, F)** Effect of two potential inhibitors of TGFβ-induced fibrosis (SB431542 and JQ1) on **(E)** IL11 secretion and **(F)** *ACTA2* and *COL1A1* expression. **(G)** αSMA and COL1A1 expression in TGFβ-treated organoids with/without indicated inhibitors. Representative images shown. * p < 0.01, ** p < 0.05, *** p < 0.001, **** p < 0.0001. *N* = 3 with each repeat comprised of 15 organoids pooled from 3 independent differentiations and treatments, One-Way ANOVA with multiple comparisons (Fisher’s LSD).

We then explored the utility of this system to test potential inhibitors of fibrosis, selecting two compounds that inhibit pathways currently under investigation in clinical trials for myeloid malignancies – SB431542, an inhibitor of the TGFβ superfamily type I activin receptors, and a bromodomain inhibitor JQ1 (55). TGFβ–induced expression of IL11 was completely inhibited by both treatments (Fig. 5E), while TGFβ–induced over-expression of *ACTA2* and *COL1A1* was normalized by JQ1 and reduced by 1.5 fold and 2.5 fold respectively by SB431542 (Fig. 5F), both at transcript and protein level (Fig. 5G). Together, these data confirm that the bone marrow organoids provide an efficient model of malignant bone marrow fibrosis and enable screening for efficacy of potential pharmacological modulators.

### Organoid ‘niche remodelling’ and fibrosis occurred following engraftment with cells from patients with myelofibrosis but not healthy donors

Having confirmed substantial homology to native bone marrow, we hypothesized that the organoids may support engraftment of primary cells from patients with hematological malignancies, enabling modelling of cancer-stroma interactions. Given the fibrosis observed following treatment with TGFβ, and the current lack of adequate *in vitro* and *in vivo* systems for modelling cancer-induced bone marrow fibrosis, we first seeded the organoids with cells from healthy donors and patients with myelofibrosis, and tested the impact of engraftment on remodelling of the bone marrow organoid “niche”.

Organoids were seeded with CD34^+^ HSPCs from 4 mobilised healthy donors and 7 patients with myelofibrosis. Donor cells were labelled with the plekstrin homology domain dye CellVue Claret, and 5000 donor cells seeded into each well of a 96-well ultra-low attachment plate containing organoids (Fig. 6A). CellVue labelling enabled the identification and tracking of donor cells within the organoid milieu. Confocal Z-stack imaging confirmed that labelled cells from healthy donors and patients with myelofibrosis efficiently engrafted and were distributed throughout the organoid architecture (Fig. 6B, Supp. Fig. 6).

**Figure 6:**
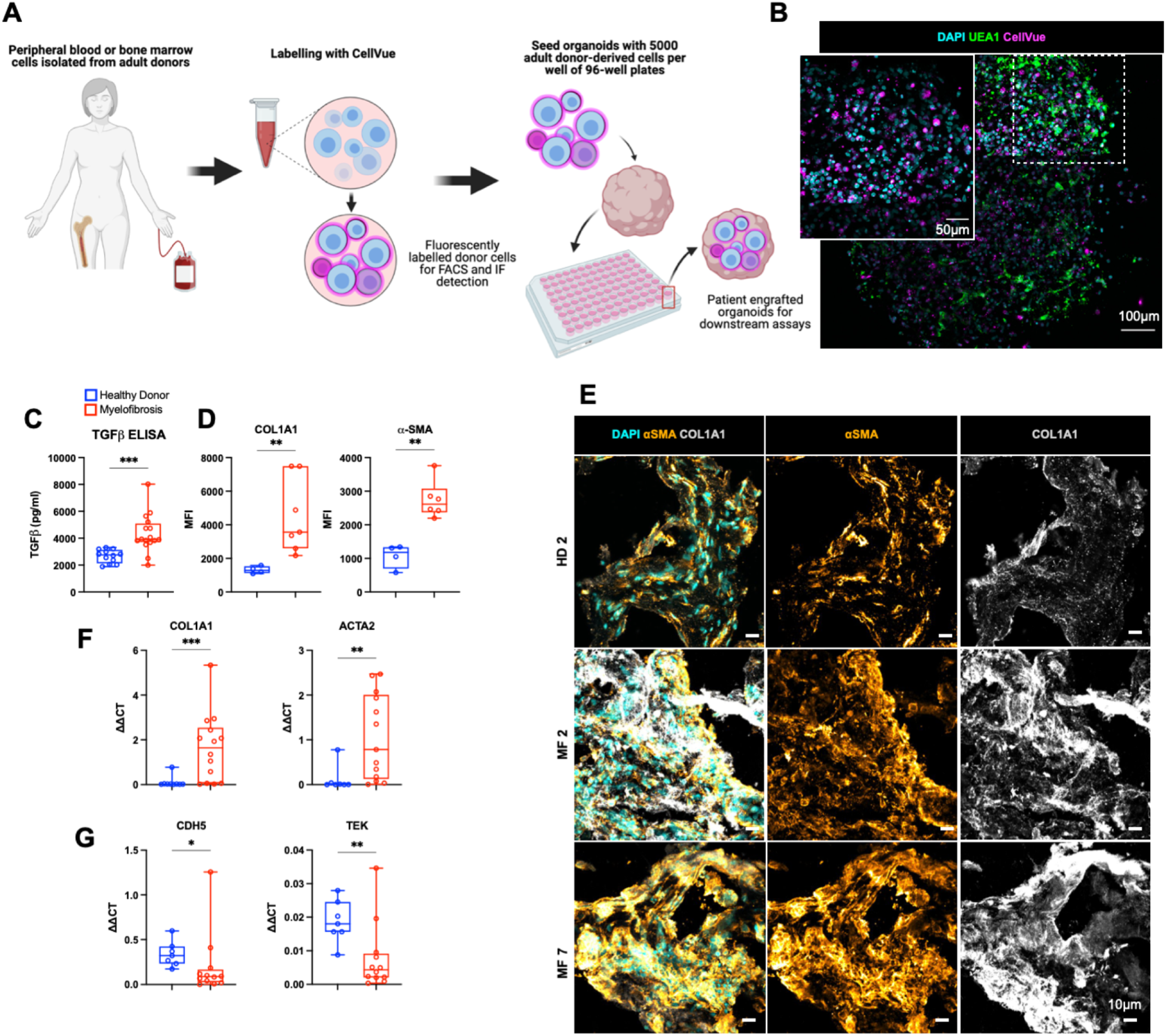
Engraftment of cells from patients with myelofibrosis, but not healthy donors, results in organoid ‘niche remodelling’ and fibrosis. **(A)** Cryopreserved peripheral blood or bone marrow cells from healthy donors and patients with blood cancers were labelled with CellVue and 5000 donor cells seeded into each well of a 96-well plate containing individual organoids. **(B)** Maximum-intensity projection of confocal Z-stack of a whole engrafted organoid 72 hours after seeding of the wells with donor cells, indicating donor cells engrafted throughout the volume of the organoids. **(C)** Soluble TGFβ in organoids engrafted by cells from myelofibrosis patients and controls. **(D, E, F,G)** Comparison of organoids engrafted with healthy donor and myelofibrosis cells for **(D)** αSMA and collagen 1 immunofluorescence staining quantification and **(E)** representative images; **(F)** *ACTA2* and *Col1A1* gene expression; **(G)** *CDH5* and *TIE2* expression. Schematic A created on Biorender.com. * p < 0.01, ** p < 0.05, *** p < 0.001, **** p < 0.0001 for Mann-Whitney test, n = *4* healthy donors; n = 7 MF samples. Representative images shown. **See also Figure S6**.

After 14 days, engrafted organoids were assessed for fibrosis. Soluble TGFβ levels were significantly elevated in the culture media of organoids engrafted with myelofibrosis cells as compared to healthy donor-engrafted samples (Fig. 6C). Immunofluorescence imaging showed a significant increase in aSMA and collagen 1 in organoids engrafted with cells from 7/7 patients with myelofibrosis, at protein level (Fig. 6D & 6E, Supp. Fig. 6B) as well as gene expression (Fig. 6F). Imaging of the organoids suggested a loss in vascular integrity in organoids engrafted with myelofibrosis cells (Suppl. Fig. 6A), therefore we investigated the expression of endothelial cell-associated genes *CDH5* and *TIE2*, and found reduced expression (Fig. 6G), confirming that engraftment of organoids with myelofibrosis cells initiated extensive niche remodelling, recapitulating the features of bone marrow biopsies from patients.

### Bone marrow organoids support the engraftment, survival and proliferation of primary cells from a range of hematological malignancies

Finally, we investigated whether primary human cells of other blood cancer types would also successfully engraft the bone marrow organoids. We focused on hematological malignancies that are particularly challenging to maintain *ex vivo* and/or model *in vivo* – infant B-cell acute lymphoblastic leukemia (iALL), chronic myeloid leukemia (CML), and multiple myeloma – to explore whether the organoids could improve survival of cells *ex vivo*, thereby enabling mechanistic studies and target screening for these cancer types.

As before, cells were labelled with CellVue Claret prior to seeding into 96-well plates containing organoids or plates containing equivalent media alone without organoids. Donor cells rapidly engrafted and were observed dispersed throughout the organoid volume (Fig. 7A, B). Multiple myeloma cells were co-stained for CD38, confirming that cells derived from the malignant plasma cell clone had successfully engrafted (Fig. 7C). We assessing the *ex vivo* viability of plasma cells that had been isolated from bone marrow aspirates of 3 patients with multiple myeloma and cryopreserved prior to thawing and engraftment in the organoids. Survival of myeloma cells was dramatically improved by seeding in the organoids. Whereas myeloma cells were less than 20% viable only 48 hours after seeding in wells with media alone, in stark contrast, the myeloma cells expanded and remained >90% viable more than 10 days after engraftment into organoids (Fig. 7D, E, F). These data confirm that the organoids provide a supportive niche for primary blood cancer cells from patients that are otherwise non-viable in ex-vivo conditions after sampling.

**Figure 7:**
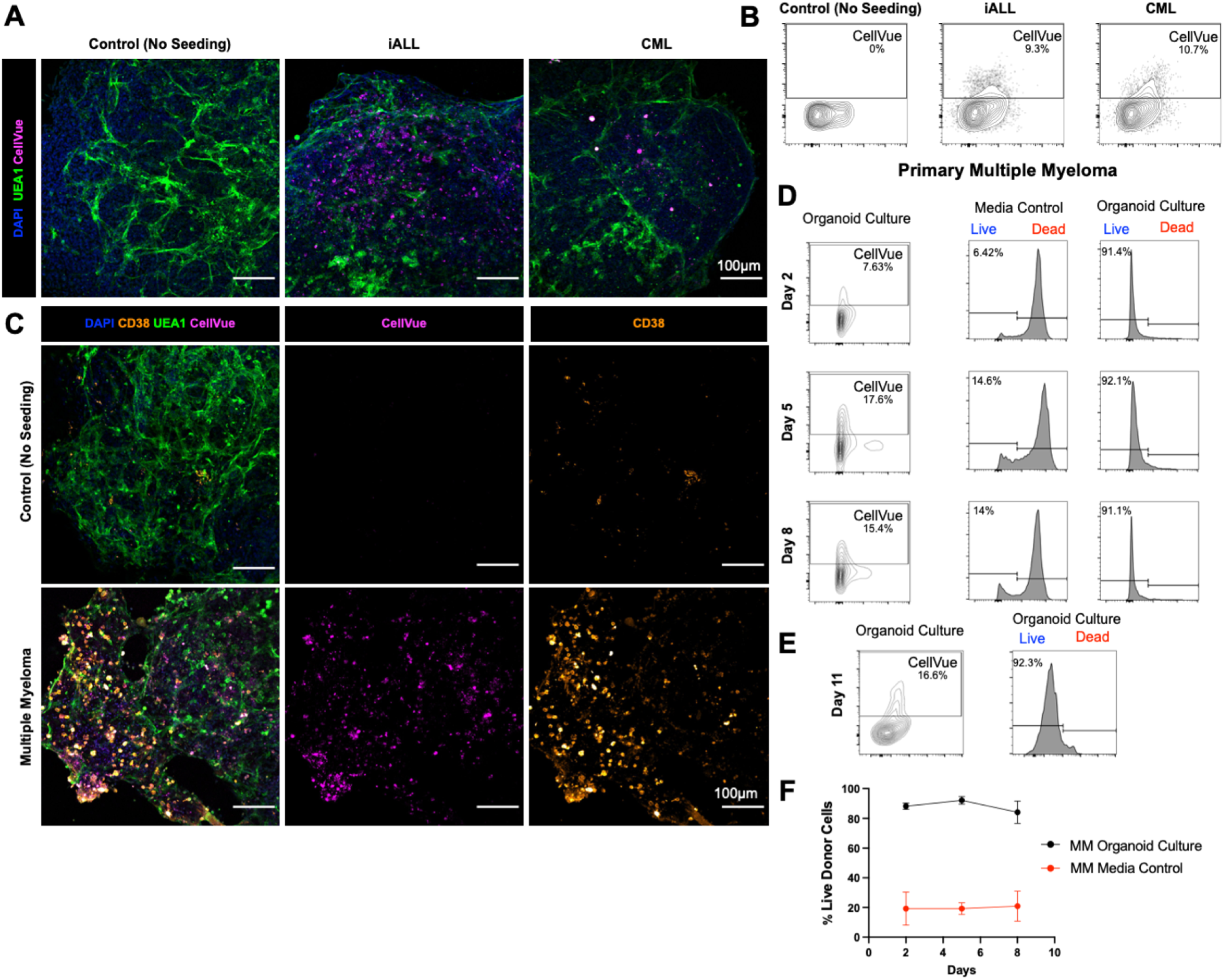
Bone marrow organoids support the engraftment, survival and proliferation of cells from patients with myeloid and lymphoid hematological malignancies. **(A, B)** Organoids engrafted with CellVue-labelled primary infant acute lymphoblastic leukemia (iALL) and chronic myeloid leukemia (CML) cells. CellVue^+^ cells engrafted throughout the volume of organoids. **(C)** Organoids seeded with CD138^+^ cells isolated from myeloma patients show CellVue^+^ CD38^+^ plasma cell engraftment. **(D)** Proportion and viability of CellVue^+^ cells within organoids at days 2, 5, and 8 after seeding. **(D)** Viability of myeloma cells within organoids vs. in wells containing media alone. **(E)** Proportion and viability of myeloma cells within organoids at day 11 (media control cells no longer viable). (F) Survival of primary multiple myeloma cells from 3 donors (and 3 independent differentiations) were assessed in media only and organoid culture conditions, with survival consistent across engraftments.

## DISCUSSION

Here we describe the development of a protocol generating vascularised bone marrow organoids that faithfully model key cellular, molecular and architectural features of myelopoietic bone marrow including stromal cells, lumen-forming vasculature and myeloid cell types. We demonstrate the utility of these organoids for modelling cancer-induced perturbations to the bone marrow niche and myelofibrosis. Treatment of organoids with TGFβ, the primary cytokine driving myelofibrosis, induced organoid fibrosis, enabling target prioritization and screening of potential inhibitors. Fibrosis also occurred following engraftment of organoids with HSPCs from patients with myelofibrosis, but not healthy donors. The ability to reliably model bone marrow fibrosis is an important advance, as the lack of adequate *in vitro* and *in vivo* models currently hampers efficient pre-clinical validation of strategies aiming to reduce or prevent fibrosis, which is a huge unmet need for patients with myeloproliferative neoplasms and other blood cancers (56). As the organoids are highly reproductible and feasible to generate at scale in 96- or 384-well plate format, this system presents an ideal platform for high-throughput target screens using pharmacological or genetic modulation.

We also show that cells from patients with both myeloid and lymphoid malignancies readily engraft and survive within the organoids, including cancer cell types that are notoriously difficult to maintain *ex vivo*. Remarkably, malignant cells from patients with multiple myeloma that had been cryopreserved and thawed prior to use were sustained by the organoids for more than 10 days, while rapidly losing viability when plated *in vitro* without organoid support. The ability to maintain survival of myeloma cells and study their interactions with a hematopoietic niche will enable pre-clinical drug testing and study of disease mechanisms using primary cells from patients, which is currently a significant obstacle to translational research in this disease.

A key limitation of current human *ex vivo* bone marrow models has been a lack of sinusoidal-like endothelium, with many systems reliant on human umbilical vein endothelial cells (HUVEC) (13). We show here that the addition of VEGFC, recently shown to support the bone marrow perivascular niche (32) drives the generation of vasculature and supporting stroma that are more specialized for hematopoietic support. The resulting bone marrow organoids thereby present a unique opportunity to study the hematopoietic-niche crosstalk that underpins healthy hematopoiesis, and how perturbations to these regulatory interactions are permissive for the emergence and progression of cancers (57).

While this system represents a significant advance, in the current iteration, no osteoid lineage, lymphoid cells, smooth muscle cells or adipocytes are generated, as the current differentiation protocol was optimized primarily to study myeloid malignancies and cancer-associated bone marrow fibrosis. However, maintenance of cells from B-cell leukemia and plasma cell malignancies provides proof of principle that the organoids can be used to support a range of bone marrow cancers and cell types, paving the way for customization to support other relevant studies. Further optimization of cytokine and growth factor cocktails, plus introduction of re-circulating flow (58), may enable future iterations of the organoids to better mimic bone marrow physiology. A protocol for ectopic implantation of human bone marrow MSC-derived ossicles has previously been shown to support engraftment of adult HSCs *in vivo* (59). Although the iPSC-derived organoids are not an *in* vivo system, they are substantially more efficient to generate than *in vivo* ossicles (weeks *vs*. months), and do not require human bone marrow or platelet lysates, which are hard to source and may induce experimental variability.

The development of organoids has been transformative in other disease settings, e.g. cerebral, lung and kidney disease modelling (60). This platform may similarly be an enabling technology for the interrogation of disease mechanisms in hematological cancers and development of novel therapies using human cells in a tissue-relevant system. Importantly, this platform is likely to reduce reliance on animal models. Target identification and screening using a species-specific, clinically relevant *ex vivo* model that can incorporate primary cells from patients may accelerate and increase the success rate of clinical translation.

## Materials and Methods

### iPSC Culture and Differentiation

A Gibco Human Episomal iPSC (Thermo Fisher Scientific Cat#A18945) line was maintained in StemFlex medium (Thermo Fisher Scientific Cat # A3349401) and on Geltrex (Thermo Fisher Scientific Cat#A1569601)-coated 6-well plates. The iPSC line was karyotyped prior to use (20). Cells were passaged as clumps using EDTA at 0.02% in PBS (0.5mM, Sigma Cat#E8008), and were freshly thawed or passaged for differentiation maintained in StemFlex supplemented with RevitaCell (Thermo Fisher Scientific Cat#2644501). Cultures were maintained at 37°C and 5% CO_2_.

Human iPSC were differentiated to mesodermal, endothelial and haematopoietic lineages in a 3-stage process as described in Figure 1A and the main manuscript text using APEL2 (StemCell Technologies Cat#05275)(61) supplemented with Bone Morphogenic Protein-4 (BMP4, Thermo Fisher Scientific Cat#PHC9531), Fibroblast Growth Factor-2 (FGF2, StemCell Technologies Cat#78134.1), Vascular Endothelial Growth Factor-A (VEGF-165, StemCell Technologies Cat#78159.1), Stem Cell Factor (hSCF, StemCell Technologies Cat#78062), recombinant Fms-like tyrosine kinase-3 (Flt3, StemCell Technologies Cat#78009), Erythropoeitin (EPO, StemCell Technologies, Cat#78007), Thrombopoeitin (TPO, StemCell Technologies, Cat#78210), Granuolocytic Colony-Stimulating Factor (G-CSF, StemCell Technologies, Cat#78012), Interleukin-3 (IL3, StemCell Technologies, Cat#78194) and Interleukin-6 (IL6, StemCell Technologies, Cat#78050).

### Immunofluorescence staining

Sections were blocked using 2% Goat Serum (Thermo Fisher Scientific, Cat#31872) 1% Bovine Serum Albumin (BSA) (Sigma, Cat#A9418) prior to primary antibody labelling with antibody diluted in 1% BSA, sequential PBS washes, and finally secondary labelling with AlexaFluor conjugates. Whole organoid blocking solution was further supplemented with Triton X100, Tween, and Sodium deoxycholate as described by Wimmer *et al* (18).

Sprouting organoids were imaged within hydrogels in 8-well microslides (Ibidi, Cat#80806), while whole organoids were labelled in 15mL Falcons before embedding in 0.5% Agarose within 8-well microslides. Whole organoids were subject to serial dehydration (50%, 70%, 90%, 100%) within microslides before clearance with Ethyl Cinnamate and subsequent imaging. Sections were prepared by embedding fixed organoids in Optimal Cutting Temperature compound (OCT, VWR Cat#361603E) before sectioning onto Poly-L-Lysine covered slides. Slides were washed in Acetone before immunofluorescence labelling.

Samples were labelled and imaged with the following primary antibodies: anti-human CD41 (BioLegend Cat# 303702, RRID:AB_314372), anti-human CD34 (BioLegend Cat# 343502, RRID:AB_1731898), polyclonal VE-Cadherin (Thermo Fisher Scientific Cat# 36-1900, RRID:AB_2533243), PDGFRB (CD140b) (Abcam, Cat# ab215978, RRID:AB_2894841), alpha-Smooth Muscle Actin (Millipore, Cat# 113200-500UL, RRID:AB_564184), CD71 (Thermo Fisher Scientific, Cat# 14-0719-82, RRID:AB_467338), and collagen Type I (Sigma-Aldrich Cat# C2456, RRID:AB_476836). Biotinylated Ulex Europaeus Agglutinin I (UEAI, Vector Laboratories, B0-1065-2) was used to label the vasculature. Secondary antibody labelling was performed using the following Alexa Fluor conjugates: Goat anti-Mouse IgG2a Cross-Adsorbed Secondary Antibody, Alexa Fluor 488 (Thermo Fisher Scientific Cat# 14-0719-82, RRID:AB_467338), Goat anti-Mouse IgG1 Cross-Adsorbed Secondary Antibody, Alexa Fluor 568 (Thermo Fisher Scientific Cat# A-21124, RRID:AB_2535766), Goat anti-Mouse IgG (H+L) Cross-Adsorbed Secondary Antibody, Alexa Fluor 488 (Thermo Fisher Scientific Cat# A-11001, RRID:AB_2534069), Streptavidin, Alexa Fluor^®^ 568 conjugate (Thermo Fisher Scientific Cat#S-11226), Goat anti-Rabbit IgG (H+L) Cross-Adsorbed Secondary Antibody, Alexa Fluor 647 (Thermo Fisher Scientific Cat# A-21244, RRID:AB_2535812). Nuclear labelling was achieved using DAPI (Thermo Fisher Scientific, Cat#62248).

### Microscopy and Image Analysis

Confocal microscopy was performed using a Zeiss LSM880 confocal AiryScan microscope with either a 25X LD LCI plan apo 0.8 NA dual immersion (420852-9871-000) or 40x C-APO NA 1.2 water immersion objective (421767-9971-711) as described previously(20). Confocal images were acquired as representative Z-stacks (with Z-resolution set to Nyquist requirements), and presented as maximum intensity projections (Fiji)(62) where stated. Histological preparations (reticulin and H&E) were imaged using a Zeiss AxioScan.Z1 slide scanner. Image analysis was performed in Fiji. For measurements of sprout radii, brightfield images acquired on an Evos (Thermo Fisher Scientific) desktop microscope. Sprout radii were measured manually by drawing and measuring a line from the centre to the tip of the sprout across 3 independent biological replicates, with between 30-50 sprouts measured per replicate.

### Histology

Organoids were fixed in neutral buffered formalin (Sigma-Aldrich, Cat#HT501128-4L) in a 15mL Falcon tube, washed twice with PBS, and then subject to serial dehydration (30%, 50%, 70%, 100%) in ethanol before immersion in Histoclear (Geneflow, Cat#A2-0101). Samples were then embedded in paraffin and sent as blocks to C&C laboratories for staining and mounting.

### Single-cell RNA-sequencing

Cells were thawed, stained with DAPI to exclude non-viable cells, and DAPI-live cells sorted on a Becton Dickinson Aria Fusion with 100nm nozzle as per recommendations in the 10x Genomics Single Cell Protocols – Cell Preparation Guide. 10,000 live cells per sample were sorted into 2μL PBS/0.05% BSA (non-acetylated) and the cell number/volume adjusted to the target for loading onto the 10x Chromium Controller. Samples were processed according to the 10x protocol using the Chromium Single Cell 3’ library and Gel Bead Kits v3.1 (10x Genomics). Cells and reagents were prepared and loaded onto the chip and into the Chromium Controller for droplet generation. Reverse transcription was conducted in the droplets and cDNA recovered through demulsification and bead purification. Pre-amplified cDNA was used for library preparation, multiplexed and sequenced on a Novaseq 6000.

### Single Cell RNA sequencing data processing and analysis

Demultiplexed FASTQ files were aligned to the human reference genome (GRCh38/hg38) using standard CellRanger (version 6.0.1) ‘cellranger count’ pipeline (10x Genomics). SingCellaR(35) (https://supatt-lab.github.io/SingCellaR.Doc/) was used for the downstream analysis. Data was first subject to quality control with the maximum percentage of mitochondrial genes, maximum detected genes and max number of UMIs set to 12%, 6,000, and 50,000, respectively. Minimum detected genes and UMIs were set to 300 and 500, respectively and genes with minimum expressing cells was set as 10. Raw expression matrix was then normalised and scaled and number of UMIs and percentage of mitochondrial reads were regressed out before a general linear model (GLM) was used to identify highly variable genes(63). Data were then subject to downstream analyses including principal component analysis (PCA), UMAP analysis (top 40 PCs were used, and n.neighbour = 120), and clustering using the Louvain method. Differentially expressed genes were calculated using ‘identifyDifferentialGenes’ function (min.log2FC = 0.3 and min.expFraction = 0.25). To compare cells from the two experimental conditions (VEGFA only and VEGFA+C), cells were downsampled so that each cell group had the same number of cells. Wilcoxon test of normalized UMIs was used to compare the gene expressions and Fisher’s exact test was used to compare the cell frequency. The resulting *P values* from both tests were combined using Fisher’s method and subsequently adjusted by Benjamini-Hochberg correction. ‘runFA2_ForceDirectedGraph’ function was used to identify the trajectories.

CellPhoneDB v 2.1.1 (https://github.com/Teichlab/cellphonedb) was performed for ligand-receptor interactions using normalized expression matrix of VEGFA +C as detailed by Garcia-Alonso *et al*. (64,65). Cell-cell interaction network between the different cell clusters from VEGFAC and Sankey plot demonstrating the interaction between TGFβ1, CXCL12, and CD44 ligands with their responding receptors from VEFGA and VEFGAC were plotted using a modified version of the CrossTalkeR R package (version 1.2.1; (66)).

We applied Symphony (39) to map cells from VEGFA+C organoids to published scRNAseq datasets from human bone marrow (36) and fetal liver and bone marrow cells (35,38) respectively. For the human bone marrow dataset, we first built the reference data using the normalized expression matrix using ‘symphony::buildReference’. For the fetal liver dataset we used the pre-built reference provided by the Symphony developer. The ‘mapQuery’ and ‘knnPredict’ function were used to map the VEGFA+C cells onto the three reference datasets.

### Flow Cytometry

Organoids were dissociated for flow cytometry analysis using collagenase Type II (Sigma Aldrich, Cat#C6885) resuspended in HEPES buffer (Sigma Aldrich, Cat#H0887) at a concentration of 20mg/mL. Samples were collected from 96-well ULA plates and collected by gravitation in a 15mL Falcon tube and subject to 2x PBS washes before resuspension in collagenase Type II. For dissociation, samples were incubated at 37°C for 5 minutes before trituration and a further 5 minute incubation. The dissociation reaction was stopped through the addition of PBS supplemented with FBS. Samples were blocked in PBS supplemented with 0.02mM EDTA and 2% FBS before antibody labelling. 10 organoids were dissociated per flow cytometry experiment.

Analysis was performed using either a Cyan Flow Cytometer (Beckman Coulter) or an Attune NxT.

Antibodies used included: Hematopoietic panel: Lineage-PE (Beckman Coulter, Cat#B29559), CD34-APC (BioLegend Cat# 343607, RRID:AB_2074356), CD11b-biotin (Streptavidin-Texas Red used as secondary) (BioLegend Cat# 305804, RRID:AB_314560), CD45 PE Cyanine 7 (BioLegend Cat# 304125, RRID:AB_10709440), CD42b APC Cyanine7 (BioLegend Cat# 303920, RRID:AB_2616778), CD41 Pacific Blue Conjugate (BioLegend Cat# 303714, RRID:AB_10696421); Stroma panel: CD71-FITC (BioLegend Cat# 334103, RRID:AB_1236432), CD140b/PDGFRB PE (BioLegend Cat# 323605, RRID:AB_2299493), Leptin Receptor -Alexa 647 (BD Biosciences Cat# 564376, RRID:AB_2738777), VCAM1 (BioLegend Cat# 305804, RRID:AB_314560), CD144 PECy7 (BioLegend Cat# 348516, RRID:AB_2687022), CD31 APC Cy7 (BioLegend Cat# 303120, RRID:AB_10640734), and CD235a (BioLegend Cat# 349107, RRID:AB_11219199). Single color stained controls and fluorescence-minus-one (FMO) controls were used for all experiments.

### Seeding of organoids with normal and malignant hematopoietic cells

Peripheral blood and bone marrow samples were collected from healthy donors and patients with hematological malignancies following provision of informed consent in accordance with the Declaration of Helsinki for sample collection, tissue banking and use in research. Peripheral blood samples from patients with myelofibrosis and healthy, mobilized apheresis donors were banked under the INForMed Study, University of Oxford (IRAS: 199833; REC 16/LO/1376). Following thawing, mononuclear cells were stained with anti-human CD34 BV650 (Biolegend, Cat# 343624, RRID:AB_2632725) for 20min at RT, washed and live cells (DAPI negative) were sorted using a Becton Dickinson FACSAria™ Fusion Cell Sorter with 100nm nozzle into 1.5ml Eppendorf tubes prior to seeding in organoids. Bone marrow aspirate samples from patients with multiple myeloma were banked under the Oxford Radcliffe Biobank (Oxford Clinical Research Ethics Committee 09/H0606/5/5, project 16/A185). Myeloma cells were selected from total bone marrow mononuclear cells using anti-CD138 magnetic bead enrichment (StemCell Technologies, Cat #17887) prior to cryopreservation and use. B-ALL cells were derived from a recently published model of iALL in which the t(4;11)/MLL-AF4 translocation was introduced into primary human fetal liver hematopoietic cells by CRISPR-Cas9 gene editing prior to transplantation into sub-lethally irradiated immunodeficient mice (67). Donated fetal tissue was provided by the Human Developmental Biology Resource (HDBR, www.hdbr.org), regulated by the UK Human Tissue Authority (HTA, www.hta.gov.uk) and covered under ethics (REC: 18/NE/0290 and 18/LO/0822). Cells were harvested from bone marrow of leukemic mice at 17-18 weeks following transplant, and total bone marrow cells cryopreserved. Following thawing and prior to seeding in the organoids, human CD45+ were selected using magnetic microbeads (Miltenyi Biotech, Cat# 130-045-801).

Cells were cryopreserved in FCS with 10% DMSO and thawed by warming briefly at 37°C, gradual dilution into RPMI-1640 supplemented with 10% FCS and 0.1mg/mL DNase I, centrifuged at 500G for 5 minutes and washed in FACS buffer (PBS + 2mM EDTA + 10% FCS). Prior to seeding, donor cells were labelled with CellVue Claret Far Red Fluorescent Cell Linker Mini Kit for General Membrane Labelling (Sigma Aldrich, Cat#MINCLARET-1KT) following manufacturer instructions. For seeding of organoids, each well of a 96-well plate containing 1-2 individual organoids or media alone were seeded with 5000 cells per well and cultured for up to 14 days in StemPro (Thermo Fisher Scientific, Cat#10639011) supplemented with Phase IV cytokines. Wells seeded with iALL cells were further supplemented with IL7. Half of the media was replaced every 2-3 days. On collection day, organoids were either fixed for imaging or digested for flow cytometry analysis/RNA extraction. The composition of the engrafted organoids was analyzed by flow cytometry using either an LSR Fortessa X50 (BD Biosciences) or an Attune NxT.

### Quantitative Real-Time Polymerase Chain Reaction (qRT-PCR)

Whole organoids were processed using either the Micro RNEasy Kit (Qiagen, Cat#74004) or Qiagen Mini RNA isolation kit (Qiagen, Cat#74104) according to the manufacturer’s instructions. Isolated RNA was quantified on the NanoDrop ND-100 (Thermo Scientific) and cDNA was prepared using using the High Capacity cDNA Reverse Transcription Kit (Applied Biosystems, Cat# 4368814) or EvoScript Universal cDNA Master (Roche, Cat#07912374001) according to the manufacturer’s instructions using standard cycling conditions. cDNA was diluted to 5ng before being combined with PowerUp SYBR Green Master Mix reagent (Applied Biosystems, Cat# A25742) and the relevant PrimeTime qRT-PCR primers (IDT), or performed using TaqMan™ Universal PCR Master Mix (Applied Biosystems) on StepOne plus machine (Applied Biosystem) (see Suppl. Table 3 for a full list of primers). The absolute expression of the respective genes was calculated using the △Ct method using *GAPDH* as an internal housekeeping control.

### Data Analysis

scRNAseq analyses were performed in R Studio (version 1.4.1106). Other statistical analyses were performed using Graph Pad PRISM 7 with statistical tests as described in relevant figure legends. *P* values defined as * p < 0.01, ** p < 0.05, *** p < 0.001, **** p < 0.0001 throughout.

## Supporting information

Supplementary material for Khan et al.

Supp Table 1 GSEA lists

## Acknowledgements

We thank all the patients who kindly donated samples and: Nawshad Hayder and Sophie Reed who helped with sample banking; the University of Birmingham TechHub Imaging Core Facilities and staff; the MRC WIMM Flow Cytometry facility; the MRC WIMM Single Cell Facility; Daniel Royston for input and advice on bone marrow architecture and histology; Zewen Kelvin Tuong for his assistance with KT; Arturs Habirovs and Ashleigh Danks for computational advice.

## Availability of data and material

Analysis scripts, raw and processed sequencing data will be made publicly available on publication of this work.

## Author Contributions

AOK conceived of and designed the differentiation protocol, BP designed fibrosis, donor cell engraftment and single cell RNASeq experiments. AOK, BP and KRM contributed to the overall design of the study and structure of the manuscript. AOK performed cell culture, differentiations, imaging experiments and subsequent analysis. AOK, MC, and JSR performed flow cytometry and qRT-PCR experiments. MC and LM performed single cell RNA sequencing experiments. RNAseq analysis was performed by AOK and GW, assisted by WXW, CM and BP. ARR and MC performed seeding and analysis of organoids with donor cells. Patient samples were banked and provided by SG, ARR, AR and BP. AOK and BP co-wrote the manuscript. All authors provided input and reviewed the manuscript.

## Consent to participate

N/A

## Consent for publication

N/A

