## Supplementary material for Khan et al. for "Human bone marrow organoids for disease modelling, discovery and validation of therapeutic targets in hematological malignancies"

#### Supplementary Figure 1 (related to Figure 1)

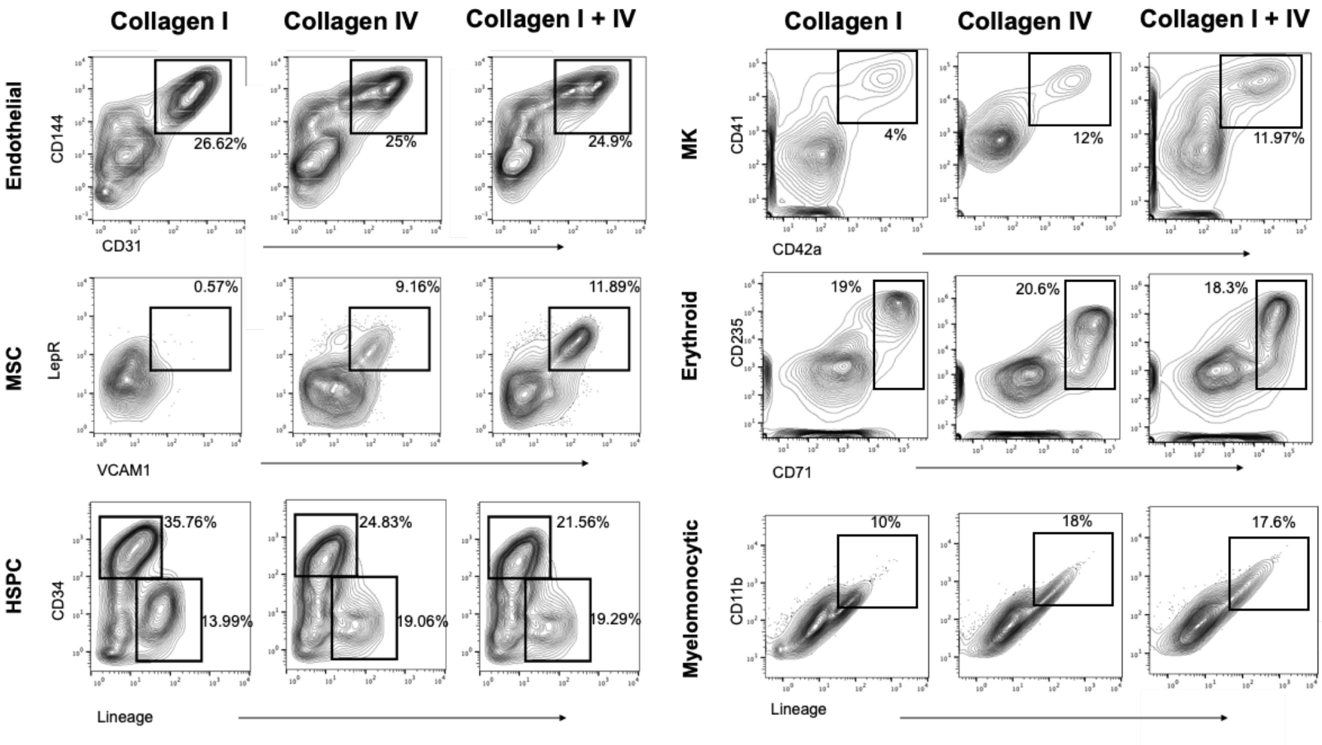


**Supplementary Figure 1, relating to Figure 1: Representative flow cytometry plots for organoids differentiated in collagen I, collagen IV and collagen I+IV Matrigel hydrogels.** Gating strategy for hematopoietic and stromal cell types is shown. Abbreviations: Hematopoietic stem and progenitor cells (HSPC); mesenchymal stromal cells (MSC), megakaryocytes (MK).

#### Supplementary Figure 2 (Related to Figure 3)

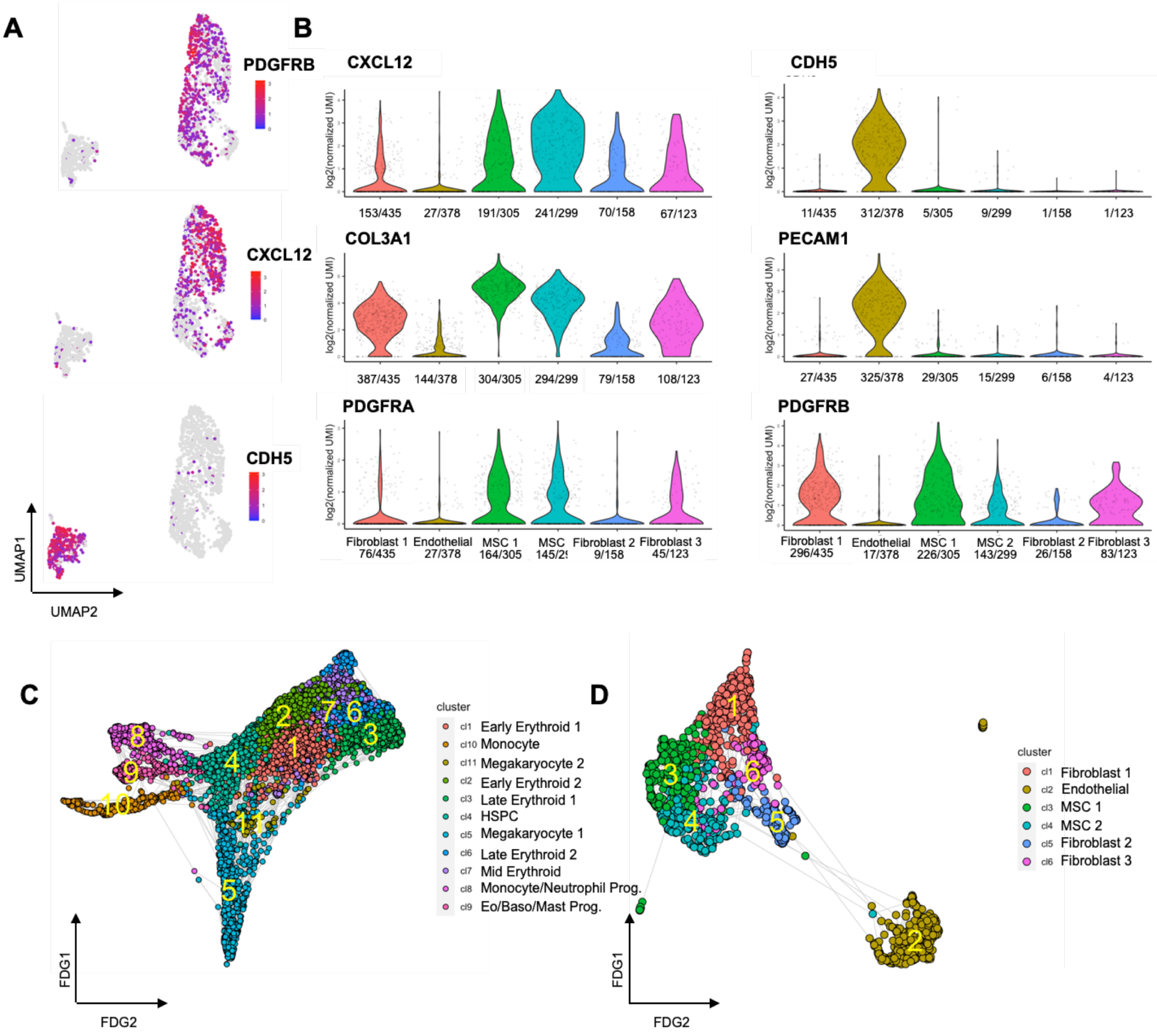


**Supplementary Figure 2, relating to Figure 3: Expression of canonical genes and differentiation trajectories in cell clusters from VEGFA+C organoids. (A)** Uniform Manifold and Approximation Projection (UMAP) plots showing expression of *PDGFRB* and *CXCL12* in fibroblasts and mesenchymal stromal cells (MSC) and *CDH5* in endothelial cell clusters of bone marrow organoids respectively. **(B)** Violin plots showing expression of key genes in stromal cell clusters. Number of cells in each cluster in which expression of gene was detected is indicated below the plot. **(C & D)** Force-directed graph (FDG) of **(C)** hematopoietic and **(D)** stromal cell compartments.

#### Supplementary Figure 3 (Related to Figure 3)

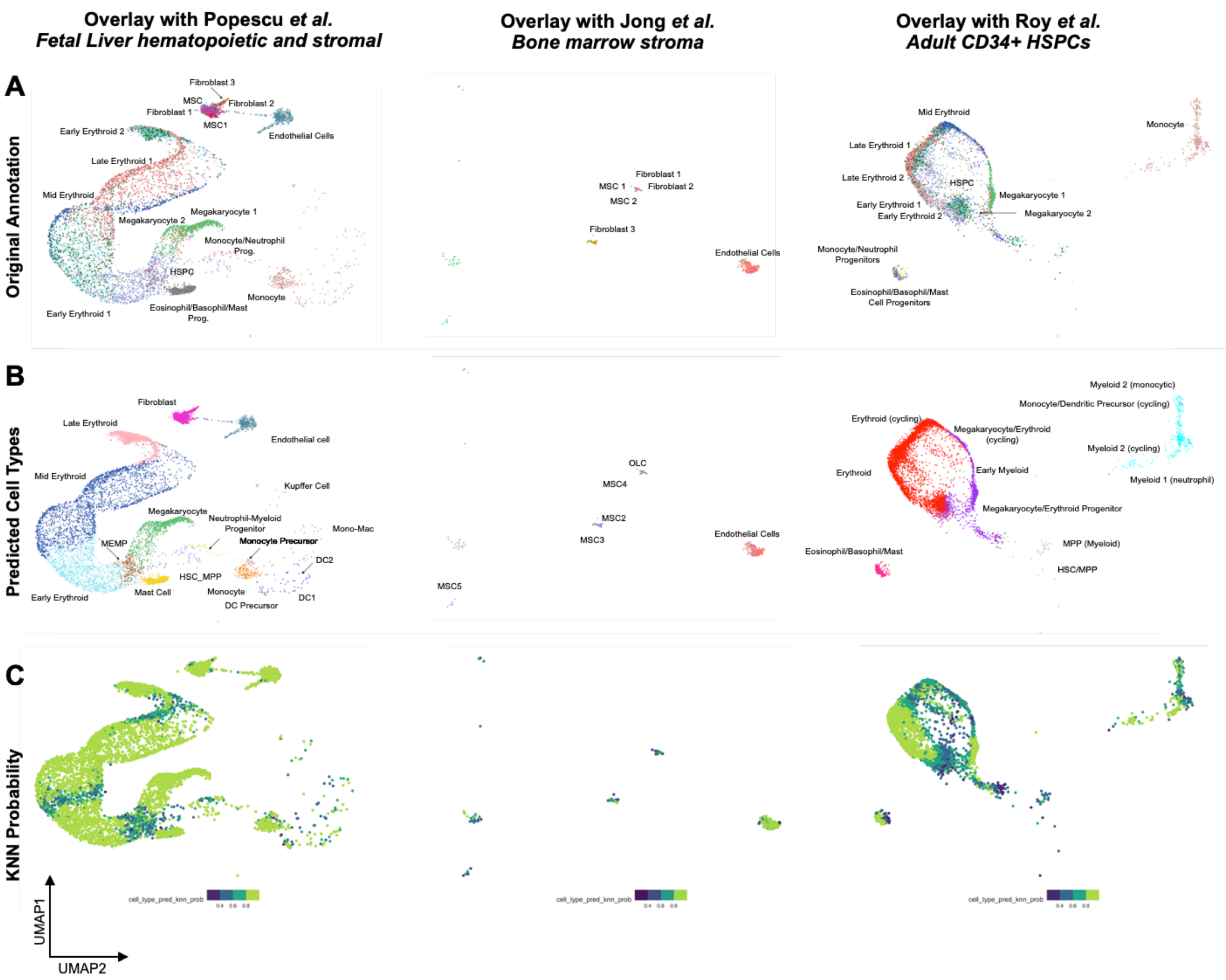


**Supplementary Figure 3, relating to Figure 3: Overlay of organoid cells to published datasets of human hematopoietic and stromal cells using the Symphony package. (A)** original annotation as per Figure 3. **(B)** Predicted cell types **(C)** KNN score, representing the spearman correlation score between the cells in the reference and query datasets, where a high score indicates good correlation between cell types.

#### Supplementary Figure 4 (Related to Figure 4)

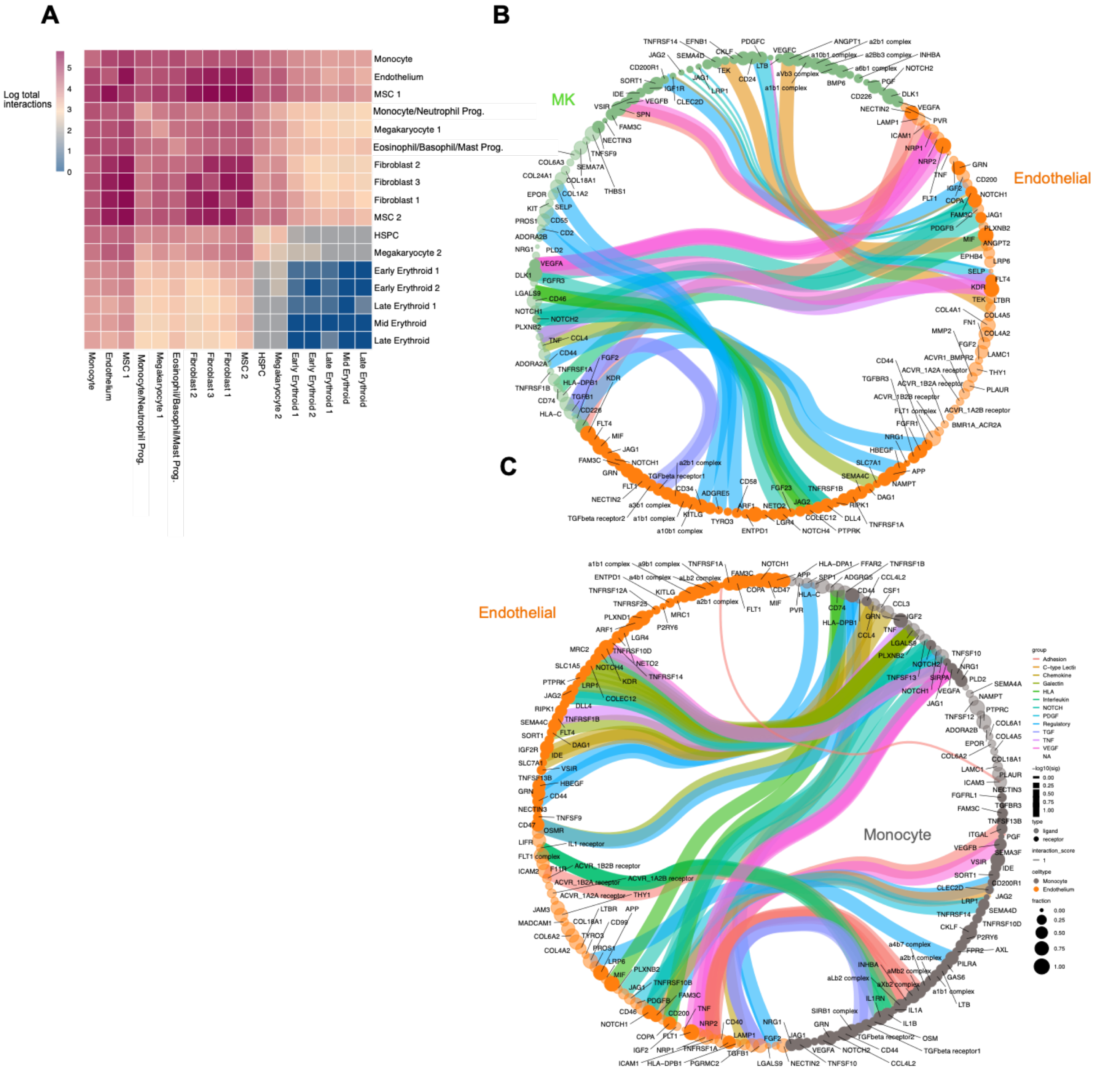


**Supplementary Figure 4, relating to Figure 4: Cell-cell interactions between hematopoietic and stromal cell types in VEGFA+C-stimulated organoids.** (**A**) Heatmap showing the log(number) of significant predicted receptor (R) – ligand (L) interactions between cell clusters identified using CellPhoneDB v2.0.1. (**B & C**) Circos plots showing interactions between **(B)** megakaryocytes and **(C)** monocytes with endothelial cells (KTPlots). Colors indicate R-L group, width of connecting band reflects log(10) p value and the size of the circle indicates the percentage of cells within the cluster that expressed the relevant receptor or ligand. Interactions were color coded by grouping, and only significant interactions were plotted.

#### Supplementary Figure 5 (Related to Figure 4)

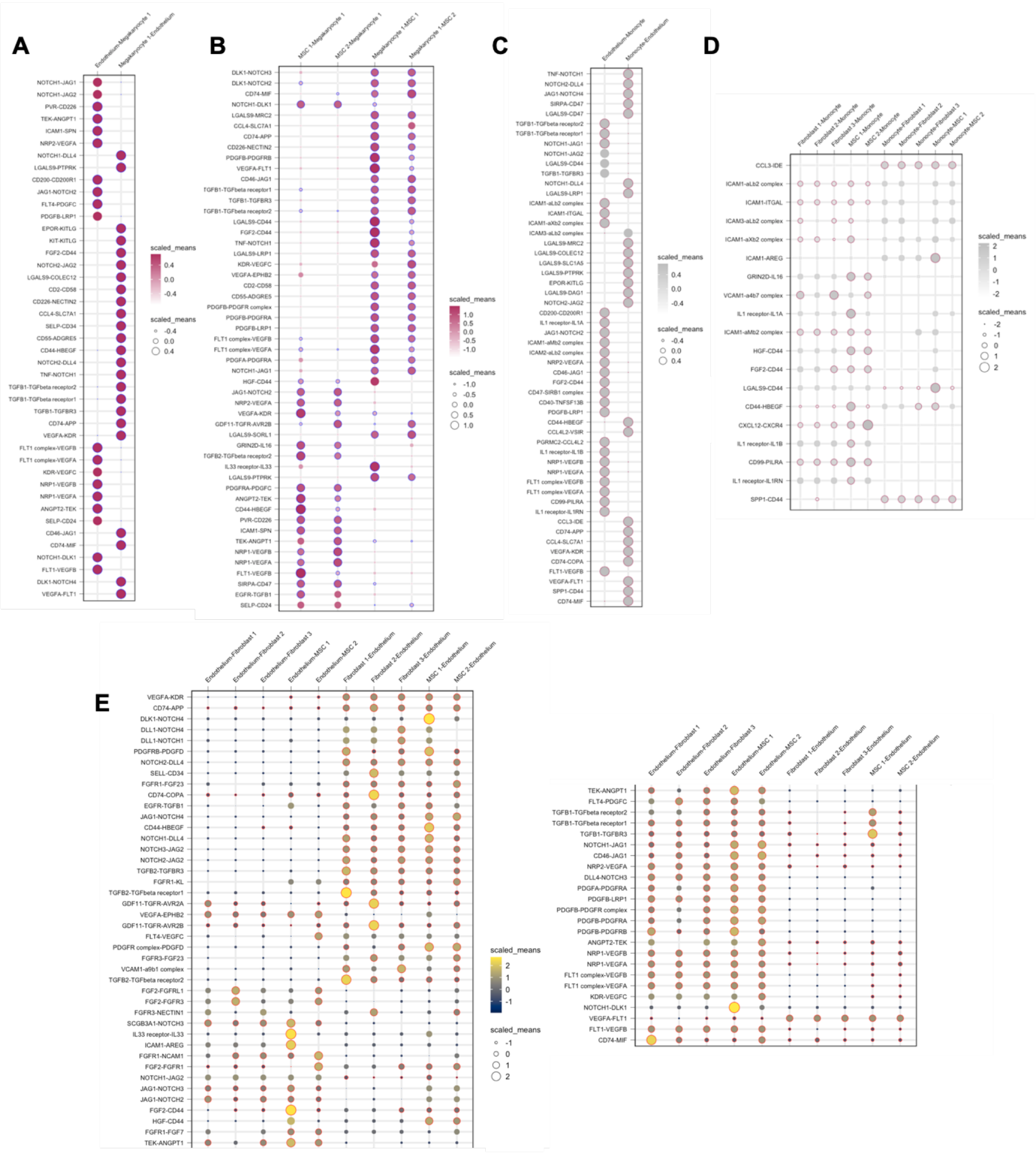


**Supplementary Figure 5, relating to Figure 4: Dot plots showing significant receptor-ligand interactions between key interacting cell types within the organoids.** Significant interactions reported by CellPhoneDB (p value < 0.05) were plotted by clusters of choice to highlight interactions of interest. Each circle is both color and size coded to indicate relevant expression level (scaled means) or the ligand:receptor pairing in the clusters of interest. **(A)** Significant endothelial cell – megakaryocyte; **(B)** mesenchymal stromal cell (MSC) - megakaryocyte; **(C)** endothelial cell – monocyte; **(D)** fibroblast/MSC – monocyte and **(E)** endothelial cell – MSC/fibroblast interacting ligand-receptor pairs.

#### Supplementary Figure 6 (Related to Figure 6)

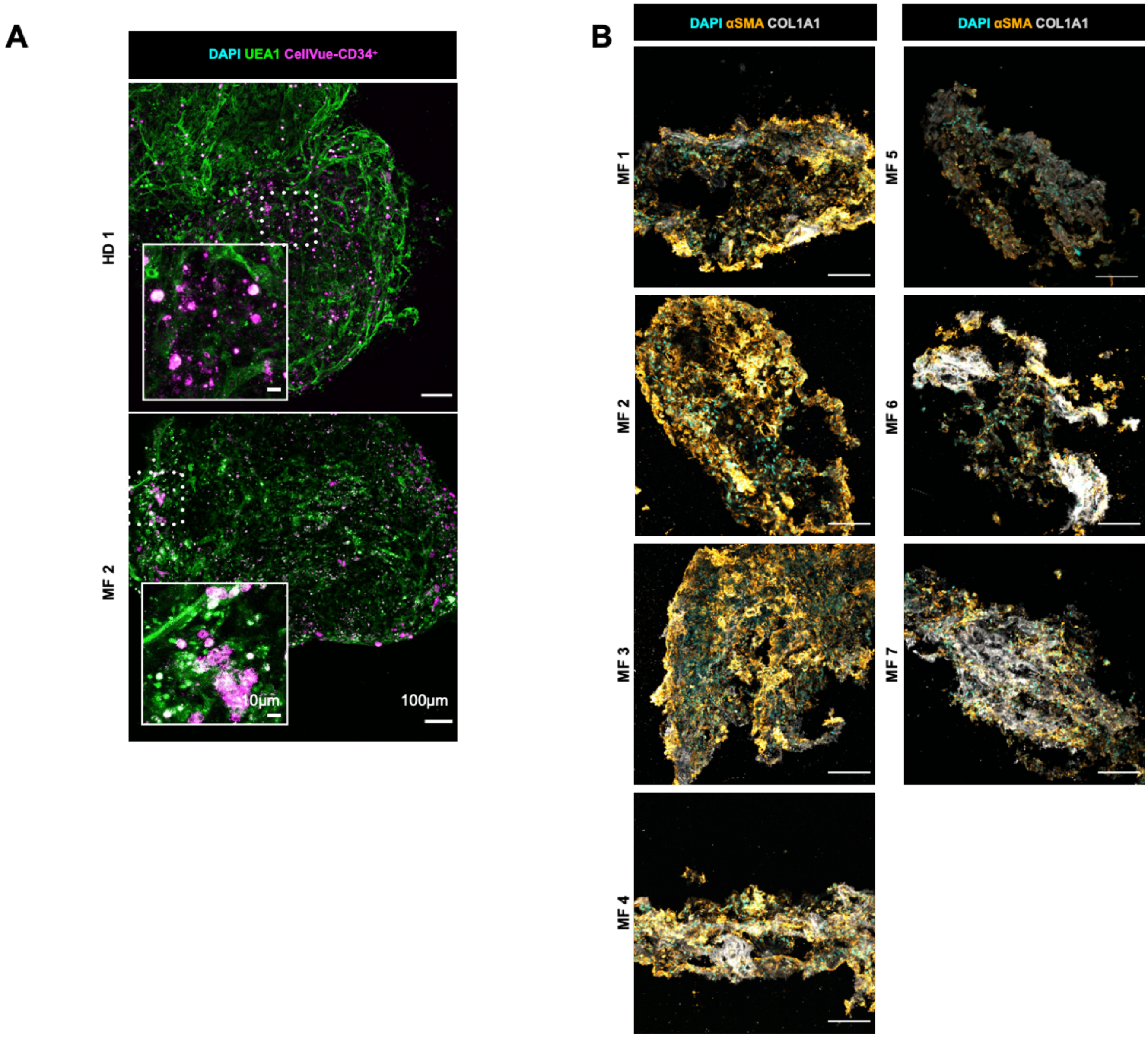


**Supplementary Figure 6, relating to Figure 6: Organoids engrafted with CD34+ hematopoietic stem/progenitor cells from healthy donors and patients with myelofibrosis. (A).** CellVue Claret-labelled healthy donor and patient cells visualized within organoids. **(B)** Increased collagen 1 deposition and αSMA expression 14 days following seeding of organoids with cells from patients with myelofibrosis.

### Supplementary Table 3 (Related to figures 5 and 6)

| Gene (target) | Primer ID |
|---|---|
| **ACTA2 (aSMA)** | Hs.PT.56a.2542642 (PrimeTime) |
| **COL1A1** | Hs.PT.58.15517795 (PrimeTime) |
| **ITGA4(VLA4)** | Hs.PT.58.40415661 (PrimeTime) |
| **FLT4 (VEGFR3)** | Hs.PT.58.19569554 (PrimeTime) |
| **VCAM1** | Hs.PT.58.20405152 (PrimeTime) |
| **FGF4** | Hs.PT.58.2180933 (PrimeTime) |
| **CXCR4** | Hs.PT.58.27595676 (PrimeTime) |
| **GAPDH** | Hs.PT.39a.22214836 (PrimeTime) |
| **GAPDH** | Hs02758991_g1 (TaqMan) |
| **ACTA2** | Hs00426835_g1 (TaqMan) |
| **COL1A1** | Hs00164004_m1 (TaqMan) |
| **CDH5** | Hs00901465_m1 (TaqMan) |
| **TEK** | Hs00945150_m1 (TaqMan) |

**Supplementary Table 3:** List of qRT PCR probes used.
